# Coordinated cell and chloroplast growth and its perturbation by chloroplast DNA replication inhibition in green algae

**DOI:** 10.64898/2026.05.06.723297

**Authors:** Veronika Kselíková, Adéla Vaňková, Sien Audoor, Bipasha Bhattacharjee, Firas Louis, Martín Mora-García, Rabinder Singh, Ana Álvarez González, Enes Göksal, Kateřina Bišová

## Abstract

Coordination among cell growth, chloroplast expansion, and organelle genome dynamics is fundamental to algal physiology, yet its regulation remains unclear. We used time-resolved single-cell analyses to examine scaling relationships among cell size, chloroplast volume, nuclear dynamics, and nucleoid organization in *Desmodesmus communis* and *Chlamydomonas reinhardtii* under normal conditions and after inhibition of chloroplast DNA replication with nalidixic acid (NAL). Under control conditions, both species showed coordinated scaling among cell, chloroplast, and nuclear size, while nucleoid dynamics were driven mainly by changes in number. NAL disrupted these relationships in a species- and time-dependent manner. In *C. reinhardtii*, prolonged treatment uncoupled chloroplast and nuclear growth from cell expansion and led to fewer, enlarged nucleoids, consistent with impaired replication. In contrast, *D. communis* largely maintained coordinated scaling, with effects mainly limited to reduced nucleoid proliferation and delayed division. Temporal analyses indicated that NAL primarily affected nucleoid replication and segregation, with secondary consequences for chloroplast growth and cell-cycle progression. These findings identify chloroplast genome dynamics as a regulatory link between organelle growth and cell division.

## Introduction

The cell cycle of eukaryotic cells consists of two components: the growth cycle and the chromosome cycle. The growth cycle involves an increase in cell size and total macromolecular content. The chromosome cycle includes DNA replication and nuclear division, resulting in the duplication and separation of genetic information between daughter cells (Mitchison, 1971; Sveiczer et al., 2004). Both cycles conclude with cell division. The situation is further complicated by the presence of at least two genetically distinct compartments in all eukaryotic cells. The nuclear compartment, which includes the nucleus, endoplasmic reticulum, Golgi body, centrioles, and various vacuoles, is accompanied by mitochondria, and in the plant lineage, also by chloroplasts. The nuclear compartment, mitochondria, and chloroplasts are all essential for cell survival. Therefore, the maintenance of their genetic material, division, and distribution among daughter cells must be coordinated. Unicellular green algae often contain only a single chloroplast, which must be divided between the two (or more) daughter cells. The chloroplast can occupy up to 50% of the cell volume; chloroplast RNA accounts for about 30% of total RNA, and about 3–15% of cellular DNA (Zachleder et al., 2016). Under phototrophic conditions, the increase in cell size strictly depends on energy production and carbon assimilation from chloroplast photosynthesis, making the chloroplast the main growth driver. The increase in chloroplast size precedes and correlates with the increase in cell size. However, the chloroplast has its own cycle, similar to the nuclear cycle, consisting of a commitment point (chloroplast division entry point) for chloroplast division, chloroplast DNA (nucleoid) replication, nucleoid division, and chloroplast division (Zachleder et al., 2016; Zachleder et al., 1995). Each event in the chloroplast cycle usually precedes the corresponding event in the nuclear cycle (Zachleder et al., 1995), with chloroplast division often overlapping with nuclear division and immediately preceding cell division.

Unicellular green algae containing a single chloroplast allow for straightforward methodological separation of chloroplast-derived growth and the chloroplast cycle from the cell cycle, simplifying the study of signaling between the chloroplast and nucleus. This is in contrast to higher plants, where signals from multiple chloroplasts within the same cell can interfere (Rea et al., 2018). Two green algae, *Desmodesmus communis* (formerly *Scenedesmus quadricauda*) and *Chlamydomonas reinhardtii*, have been used as models for studying cell cycle regulation and coordination between chloroplast and cell cycles (Bišová and Zachleder, 2014; Kabeya and Miyagishima, 2013; Zachleder et al., 2016). Both divide by multiple fission, a mechanism in which the cell grows to many times its original volume and can divide into 2, 4, 8, 16, or more daughter cells; generally, 2n daughter cells are formed (Bišová and Zachleder, 2014). The number of daughter cells depends on the growth rate, and the production of multiple daughter cells allows for the maintenance of a stable daughter cell size (Bišová and Zachleder, 2014; Cross and Umen, 2015; Zachleder et al., 2016).

Nucleoids are complexes of chloroplast DNA (cpDNA) with RNA and protein (Nishimura, 2024). They are considered the functional units for various processes, including DNA replication, repair, recombination, inheritance, and transcription (Kamimura et al., 2018). Chloroplast genomes are highly polyploid, with tens to hundreds of genome copies per organelle and substantial heterogeneity in copy number between nucleoids (Nishimura, 2024; Zachleder and Cepák, 1987a). CpDNA replicates under continuous light conditions in correlation with increases in chloroplast and cell size to maintain proper DNA content per chloroplast or cell volume (Zachleder et al., 2016); this replication is independent of the timing of chloroplast division and the cell cycle (Kabeya and Miyagishima, 2013; Zachleder et al., 1995). However, the process precedes and matches the number of nuclear DNA replications (and the corresponding nuclear and cell divisions) (Zachleder et al., 2016). CpDNA synthesis can be specifically inhibited by gyrase inhibitors, such as nalidixic acid (NAL) (Robreau and Le Gal, 1974; Zachleder et al., 2004) or novobiocin (Odahara et al., 2016). The inhibitor effect on cell cycle progression in synchronized cultures was established in *D. communis* (Zachleder et al., 2004) and in *C. reinhardtii* (Kabeya and Miyagishima, 2013; Voigt and Münzner, 1989), as well as in unsynchronized cultures of *C. reinhardtii* (Odahara et al., 2016). When NAL was applied to synchronized cultures of *D. communis*, the only observed effect in the first cell cycle was a slowdown in growth, probably due to reduced accumulation of chloroplast rRNA and ribosomes (Zachleder et al., 2004). The chloroplasts divided normally in both timing and extent, but the resulting daughter cells contained fewer nucleoids (Zachleder et al., 2004). This suggested that there is no checkpoint connection between cpDNA replication and chloroplast division. This may be because cpDNA replication is cell cycle independent, while chloroplast division is cell cycle regulated by the expression of mostly nuclear-encoded chloroplast division proteins (Miyagishima et al., 2012). Under natural conditions, no such relationship is required, as the number of nucleoids per chloroplast ranges from 30 in low light to more than 100 in high light (Zachleder and Cepák, 1987a), always exceeding the number of chloroplasts formed (Nishimura, 2024). It is intriguing to ask what will happen if incubation in NAL is prolonged. In principle, there are two options: 1) cpDNA replication and division are truly independent, leading to gradual accumulation of cells lacking nucleoids in their chloroplasts, or 2) a mechanism prevents further chloroplast division once the number of nucleoids per chloroplast decreases below a certain threshold. If so, there will be sequential accumulation of cells with a certain number of nucleoids per chloroplast, and further division beyond that will be blocked. Furthermore, decreasing the amount of genetic material per chloroplast will secondarily affect chloroplast growth, as the number of chloroplast ribosomes may be decreased or limited.

Here, we compared cell and chloroplast growth, as well as the formation and division of nucleoids, in two green algae, *C. reinhardtii* and *D. communis*, grown in the presence of the DNA gyrase inhibitor NAL (Fig. 1). We show that NAL-induced replication stress selectively destabilized chloroplast nucleoid organization while largely preserving cell and organelle growth. The disruption of nucleoid dynamics was more pronounced in *C. reinhardtii*, where it led to nucleoid aggregation and consequently to cell cycle arrest, even as growth continued. This uncoupling of growth and division provides new insight into the mechanisms that coordinate growth and genome organization within organelles.

**Figure 1.**
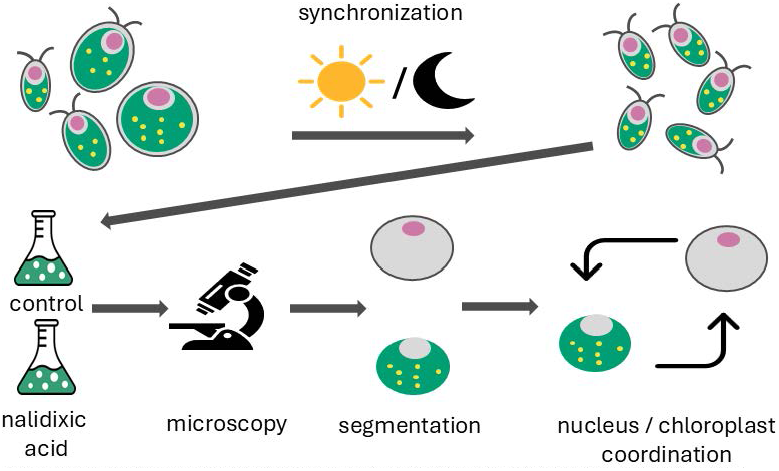
Experimental workflow. Schematic overview of the experimental design and analysis pipeline. Algal cultures were synchronized using light/dark cycles and grown under control or nalidixic acid (NAL) treatment conditions. Cells were imaged over time using microscopy, followed by segmentation and quantification of cellular and subcellular features, including cell area, chloroplast volume, nuclear area, and chloroplast nucleoid parameters (area and number). The workflow enables time-resolved analysis of coordination between cell growth, chloroplast expansion, and organelle genome dynamics, and their perturbation under inhibition of chloroplast DNA replication.

## Materials and Methods

### Organisms, culture growth conditions

*Chlamydomonas reinhardtii* strain 21gr (CC-1690; Chlamydomonas Genetics Center, Duke University, Durham, USA) and the chlorococcal alga *Desmodesmus communis* (strain no. 463, CCALA, Czech Academy of Sciences, Třeboň, Czech Republic, formerly classified as *Scenedesmus quadricauda* (Turpin) Brébisson, strain Greifswald 15) were cultivated in glass cylinders with an inner diameter of 3 cm and a volume of 300 ml. The cylinders were placed in a temperature-controlled water bath and illuminated from one side by fluorescent tubes (Osram Dulux L55W/950 daylight). Prior to the experiment, cell cultures were synchronized using alternating light and dark regimes for at least three cycles. A 12/12 cycle was used for *C. reinhardtii*, and a 14/10 cycle for *D. communis*. During synchronization, growth conditions were 30 °C, bubbling with 2% (v/v) CO2 in air, and an incident light intensity of photosynthetically active radiation at 500 μmol m^−2^ s^−1^. *C. reinhardtii* cultures were grown in modified HS medium, and *D. communis* in ½ SS medium as described by (Hlavová et al., 2016). The experiment began by diluting the daughter cells at the end of the dark period to approximately 1 × 10^6^ ml^−1^. The diluted cells were placed under light in planar glass parallel cultivation cuvettes with a light path of 2.5 cm and an inner volume of 2 liters. The growth conditions during the experiments were the same as during synchronization, except no light/dark alternation was applied. Nalidixic was added immediately after culture dilution at a concentration of 50 µg ml^−1^ for *Chlamydomonas reinhardtii* and 150 µg ml^−1^ for *D. communis*. The cultures were sampled every two hours. All experiments were conducted with at least six independent replicates.

### 4’,6-diamidine-2’-phenylindole dihydrochloride (DAPI) staining

DNA was stained with the fluorochrome 4’,6-diamidine-2’-phenylindole dihydrochloride (DAPI). *D. communis* cells were prepared using the method described by (Zachleder and Cepák, 1987b) and (Hlavová et al., 2016). One microliter of DAPI solution (5 μg/ml in 0.25 % (w/v) sucrose, 1 mM EDTA, 0.6 mM spermidine, 0.05 % (v/v) mercaptoethanol, 10 mM Tris–HCl, pH 7.6) was added to two microliters of freshly thawed *D. communis* cells and, kept for 5–10 minutes in the dark at room temperature before imaging. For *C. reinhardtii*, cells were fixed by washing in glutaraldehyde, kept at 4°C and then stained with DAPI in the same way as *D. communis*.

### Imaging and image analysis

Cells were observed using a Zeiss Axio Observer 7 microscope. For DAPI detection, 335–383 nm excitation and 420–470 nm emission filters were used. Chlorophyll was detected using 625–655 nm excitation and 665–715 nm emission filters. DAPI-stained cells were imaged as a Z-stack covering the entire cell for fluorescence microscopy. Cells were segmented in ImageJ (Schindelin et al., 2012) using the Cellpose algorithm (Stringer et al., 2021). After automatic segmentation, the results were manually curated. Cell area was derived from the segmented objects. Chlorophyll autofluorescence, used as a proxy for chloroplast size, was calculated across the entire Z-stack. Nucleoids and nuclei were identified and quantified in the maximum Z-projection of the DAPI signal using specific thresholding.

## Results

### Chloroplast and cell size scale in control cells

The algal cells began to grow immediately after exposure to light and increased in size over the course of light cultivation. This was observed in both *D. communis* (Fig. 2) and *C. reinhardtii* (Fig. 3). In both organisms, there was a strong positive scaling relationship between cell growth and chloroplast growth. Chloroplast volume increased approximately linearly with cell size (Fig. 4A, 5A), indicating coordinated growth of the chloroplast and the cell. Similarly, nuclear size scaled positively with both cell size and chloroplast volume (Fig. 4C, 5C), indicating coordinated growth of the nucleus with the cell and chloroplast. The ratio of chloroplast volume increased slightly in both species, suggesting that the chloroplast grows more than the cell (Fig. 4B, 5B). Both *D. communis* and *C. reinhardtii* divide by multiple fission, but the patterns differ. In *C. reinhardtii*, there is a prolonged G1 phase during which the cells reach a critical size, allowing them to attain commitment point and enter the cell cycle. The G1 phase is followed by alternating rounds of S and M phases and then cell division (Cross and Umen, 2015). Thus, nuclear dynamics remain unchanged until the very end of the cell cycle. In contrast, *D. communis* divides its nuclei early in the G1 phase shortly after CP attainment: first, the cells divide their nuclei into two (at about 9-12 h into light) (Fig. 2C), then into four (also at about 11-14 h into light) (Fig. 2D), and finally into eight, which is shortly followed by cell division (Fig. 2E) (Bišová and Zachleder, 2014; Zachleder et al., 2016). As a result, the cells are multinucleate for most of the cell cycle. The nuclear area reflects species-specific behavior, but in both organisms, it scaled positively with both cell size (Fig. 4C, 5C) and chloroplast volume (Fig. 4D, 5D), suggesting a close coupling to cell growth and possibly to chloroplast growth as well. The two types of nucleoid parameters, area and number, behaved differently. Nucleoid area decreased slightly with increasing cell size (Fig. 4E, 5E) and showed a weak negative trend with chloroplast size (Fig. 4F, 5F). In contrast, nucleoid number increased with both cell size (Fig. 4G, 5G) and chloroplast volume (Fig. 4H, 5H). This indicates that nucleoid proliferation is primarily driven by number rather than size.

**Figure 2.**
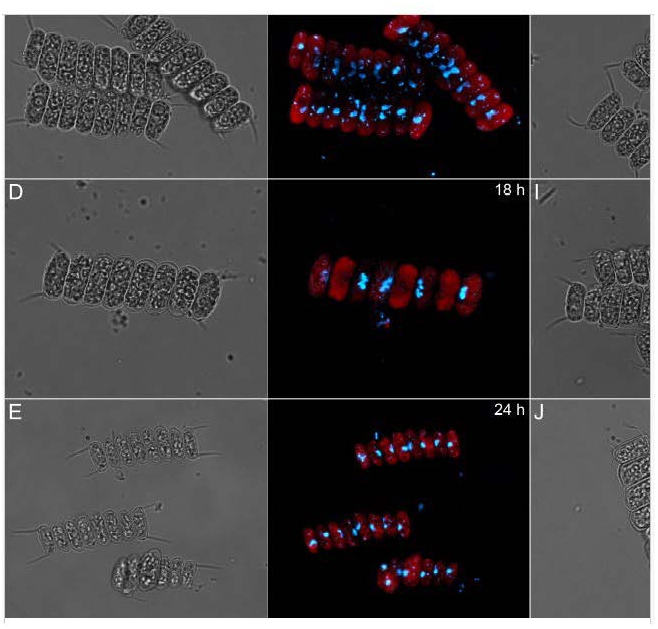
Growth and cell cycle organization and its perturbation by nalidixic acid in *Desmodesmus communis*. Growth and division patterns of control (A-E) and nalidixic acid treated (F-N) cell of *Desmodesmus communis* at different times after light switch on: 0 h (A, F), 6 h (B, G), 12 h (C, H), 18 h (D, I), 24 h (E, J), 30 h (K), 36 h (L), 42 h (M), 48 h (N), left panel – light microscopy, right panel – fluorescence microscopy after DAPI staining, bar 20 µm.

**Figure 3.**
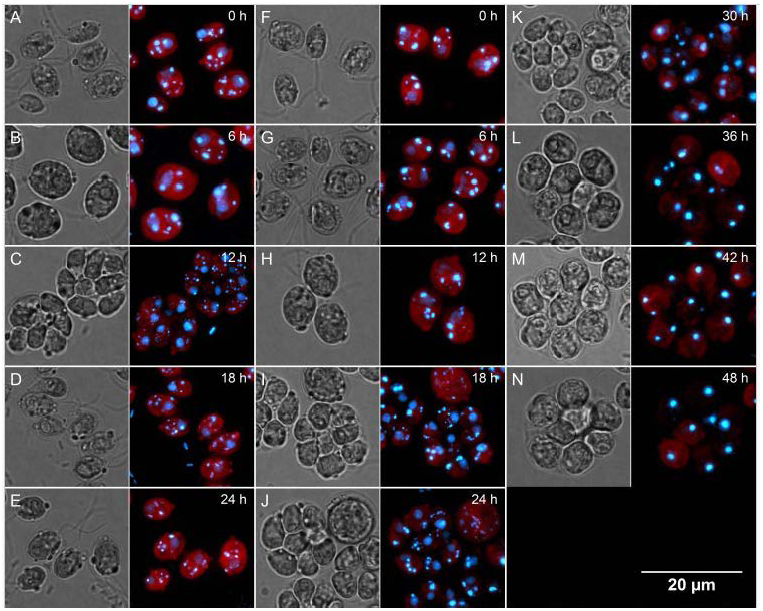
Growth and cell cycle organization and its perturbation by nalidixic acid in *Chlamydomonas reinhardtii*. Growth and division patterns of control (A-E) and nalidixic acid treated (F-N) cell of *Chlamydomonas reinhardtii* at different times after light switch on: 0 h (A, F), 6 h (B, G), 12 h (C, H), 18 h (D, I), 24 h (E, J), 30 h (K), 36 h (L), 42 h (M), 48 h (N), left panel – light microscopy, right panel – fluorescence microscopy after DAPI staining, bar 20 µm.

**Figure 4.**
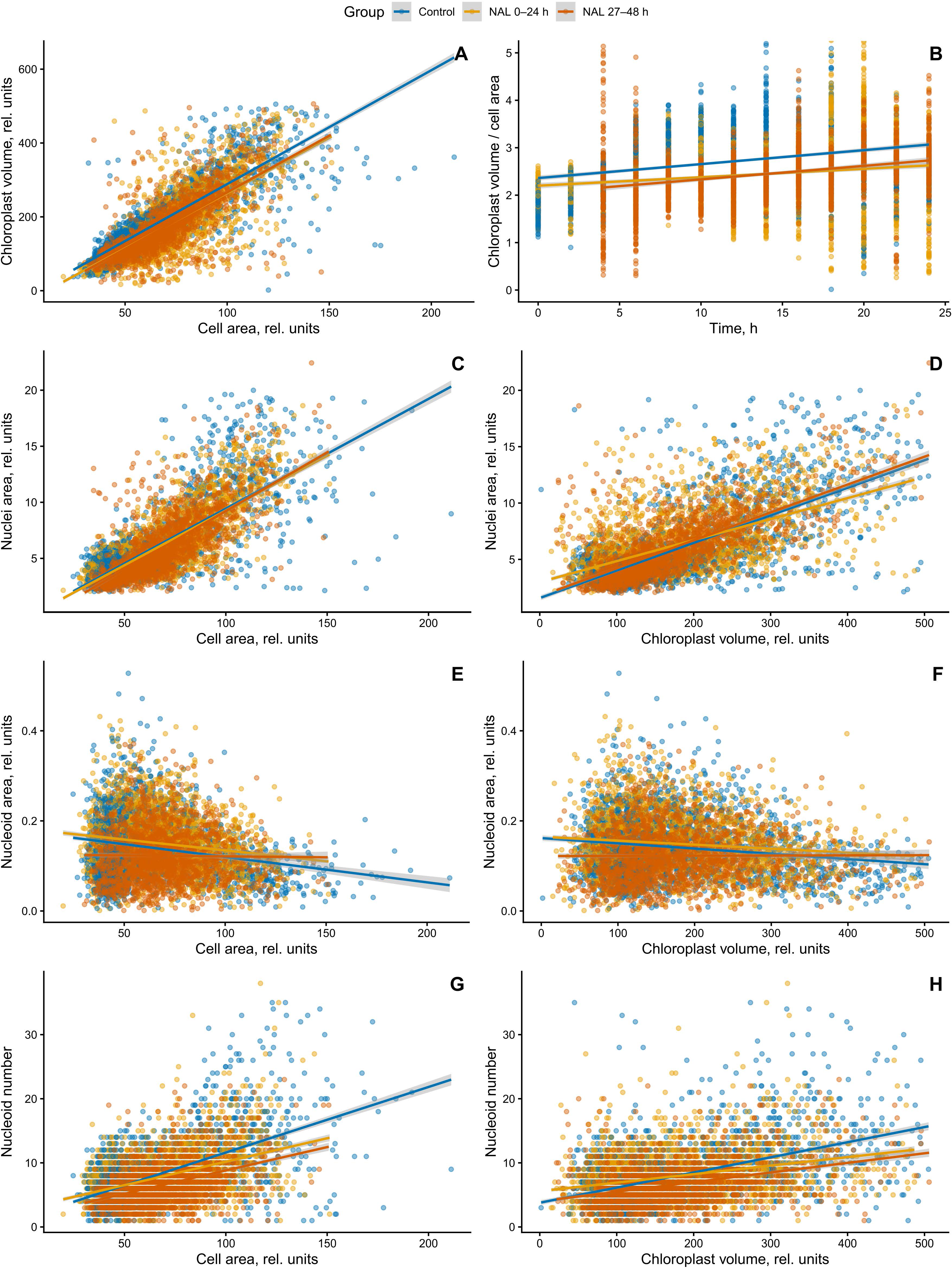
Scaling relationships between cell size, chloroplast growth, and nucleoid dynamics under control and nalidixic acid treatment in *Desmodesmus communis*. Relationships between cellular and subcellular parameters across control (blue), nalidixic acid (NAL) 0–24 h (yellow), and NAL 27–48 h (orange) conditions. (A) chloroplast volume dependency on cell area, (B) chloroplast-to-cell size ratio over time, (C, D) nuclear area dependency on cell area (C) and chloroplast volume (D), (E, F) Nucleoid area dependency on cell area (E) and chloroplast volume (F), largely maintained across conditions, although variability increases under treatment.

**Figure 5.**
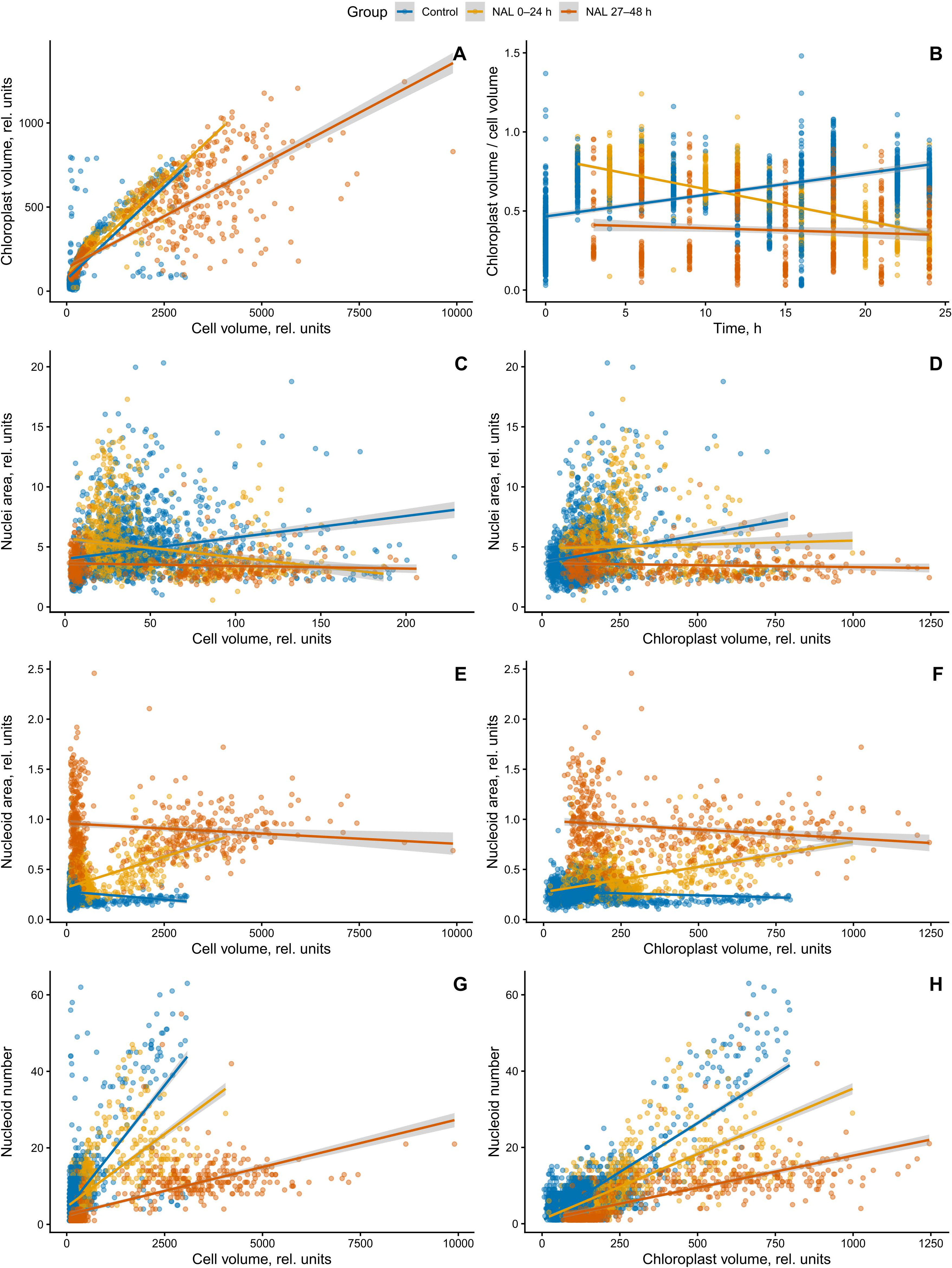
Scaling relationships between cell size, chloroplast growth, and nucleoid dynamics under control and nalidixic acid treatment in *Chlamydomonas reinhardtii*. Relationships between cellular and subcellular parameters across control (blue), nalidixic acid (NAL) 0–24 h (yellow), and NAL 27–48 h (orange) conditions. (A) chloroplast volume dependency on cell area, (B) chloroplast-to-cell size ratio over time, (C, D) nuclear area dependency on cell area (C) and chloroplast volume (D), (E, F) Nucleoid area dependency on cell area (E) and chloroplast volume (F), largely maintained across conditions, although variability increases under treatment.

### Nalidixic acid has very little effect during the first cell cycle

The first 24 hours of treatment with nalidixic acid caused very little change in chloroplast size. Chloroplast volume continued to increase with cell size, although the slope was reduced compared to control cells (Fig. 4A). This was also reflected in the chloroplast-to-cell volume (cell area) ratio. In *D. communis*, the ratio was lower than in control cells but maintained the same dynamics. In *C. reinhardtii*, the ratio decreased over time, which was in striking contrast to the increase observed in control cells (Fig. 5A). In *D. communis*, the nuclear area continued to scale positively with both cell size and chloroplast volume across all conditions (Fig. 4C, D). In *C. reinhardtii*, the slope decreased and the nuclear area became independent of both cell and chloroplast size (Fig. 5C, D).

### Prolonged treatment with nalidixic acid perturbs nucleoid dynamics and chloroplast cell size

When NAL application was extended for another 24 hours, the two species responded differently. The cells of *D. communis* reacted similarly to the early phase of treatment. In contrast, the cells of *C. reinhardtii* became more perturbed. In *D. communis*, chloroplast size scaled with cell size, and the correlation was only slightly lower than in the other two variants (Fig. 4A). Similarly, the chloroplast-to-cell size ratio was comparable to that of the NAL-treated culture in the first 24 hours (Fig. 4B), implying that while there is a slight perturbation of chloroplast size, it is not dependent on treatment duration. The scaling of nuclear area to both cell area and chloroplast volume remained unaffected and comparable to the other two variants (Fig. 4C, D). Likewise, the nucleoid area remained stable and comparable to the other two variants (Fig. 4E, F). In contrast, nucleoid numbers decreased in relation to both cell and chloroplast size (Fig. 4G, H). Although they followed the same trend as the early NAL-treated cultures, the correlation between the parameters was further weakened.

In *C. reinhardtii*, the correlation between cell size and chloroplast size decreased as the chloroplast grew less than the cell (Fig. 5A). The chloroplast-to-cell size ratio started at the low value reached by the end of the first 24 hours and remained there (Fig. 5B), suggesting that although the chloroplast made up a smaller proportion of the cell, that proportion was maintained. The nuclear area remained independent of cell and chloroplast size (Fig. 5C, D), implying that once the chloroplast proportion in the cell decreases below a certain threshold, nuclear division does not occur despite cell growth. Nucleoid area was larger than in control cells and cells treated for only 24 hours, and it became independent of both cell and chloroplast sizes (Fig. 5E, F). In contrast, nucleoid numbers still scaled with both cell and chloroplast size (Fig. 5G, H), although the correlation between them was much weaker than in the other two variants.

### Temporal dynamics of response to nalidixic acid

#### Desmodesmus communis

In general, *D. communis* growth and division were less affected by the application of NAL. The control cells coordinated their growth so that cell area (Fig. 6A), chloroplast volume (Fig. 6B), and nuclear area (Fig. 6C) increased in parallel, followed by an abrupt change at the time of cell division around 20–24 h. In the presence of NAL, growth coordination was maintained, but cell division was delayed and less synchronized (Fig. 6A, B, C). The nucleoid dynamics were more affected. In controls, nucleoid area (Fig. 6D) and count (Fig. 6E) progressively rose during growth and then dropped sharply at cell division. In treated cells, nucleoid counts (Fig. 6E) did not increase, and nucleoid area (Fig. 6D) remained higher than in controls until about 22 h. From 28 to 48 h, there was no difference, and both nucleoid counts and area remained constant despite continued cell and chloroplast growth. The application of NAL thus mainly affected nucleoid replication and segregation efficiency and secondarily slightly decreased the growth of both cells and chloroplasts.

**Figure 6.**
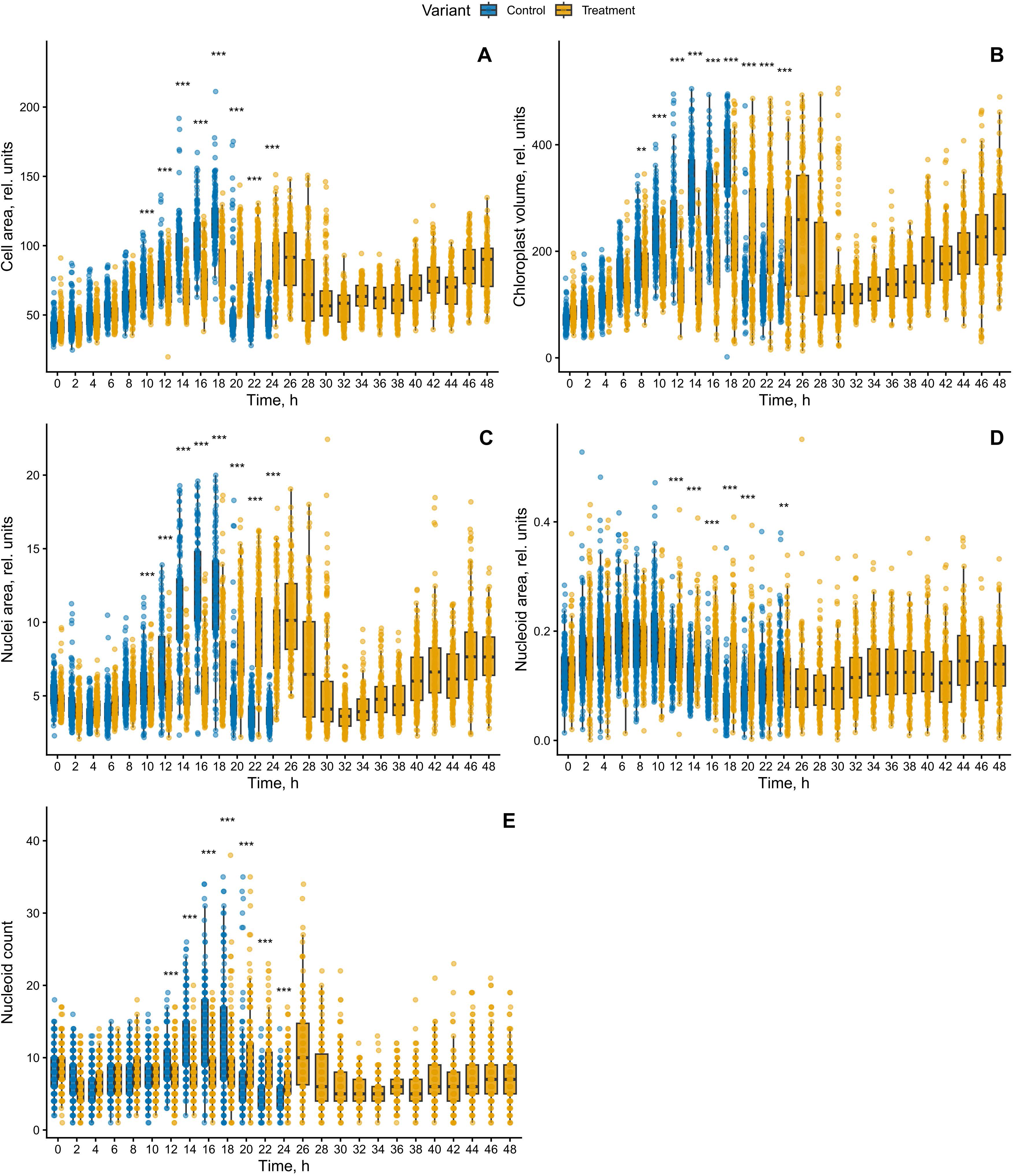
Single-cell dynamics of cellular and nucleoid parameters under control and nalidixic acid treatment in *Desmodesmus communis*. Time-course analysis of single-cell measurements in control (blue) and nalidixic acid–treated (yellow) cultures over 0–48 h, data presented as boxplots with individual cell measurements overlaid, (A) cell area, (B) chloroplast volume, (C) nuclear area, (D) chloroplast nucleoid area, and (E) nucleoid count

#### Chlamydomonas reinhardtii

Growth, division, and nucleoid dynamics were more perturbed in *C. reinhardtii*. Cell area increased progressively over time in both conditions (Fig. 7A). In control cells, there was a sharp decrease after 18 h, once the cells divided. In contrast, treated cells divided later, but the resulting daughter cells never separated and remained in palmelloid clusters formed by the four or eight sister cells (Fig. 3I-N). The cells exhibited a broader distribution and higher upper range at later time points, consistent with more heterogeneous growth of the clusters (Fig. 7A). A similar trend was observed for chloroplast volume (Fig. 7B), where treated cells showed greater heterogeneity both within and between individual samples. Nuclear area in control cells displayed an early increase followed by a reduction from around 14 h, consistent with completion of nuclear and then cell division (Fig. 7C). In treated cells, the pattern was similar but delayed and more spread out due to partial loss of cell synchrony. Once the nuclear area of treated cells stabilized, around 24 h, it remained unchanged until the end of the experiment. In control cells, nucleoid area remained virtually the same throughout the experiment (Fig. 7D), and only nucleoid numbers increased and then dropped during cell division around 14–16 h. Nucleoid counts (Fig. 7E) in treated cells increased around 16–24 h, at the time of cell and chloroplast division. Around 27 h, shortly after the clusters formed, nucleoid numbers again dropped and remained unchanged until the end of the experiment. Most striking was the behavior of nucleoid area in treated cells (Fig. 7D). It began to increase from around 14 h together with nucleoid counts. When nucleoid numbers dropped around 27 h, nucleoid area abruptly increased, reflecting the aggregation of nucleoids into fewer, larger ones. NAL thus had similar effects to *D. communis*. Yet, in *C. reinhardtii*, chloroplast growth and nucleoid dynamics were more perturbed.

**Figure 7.**
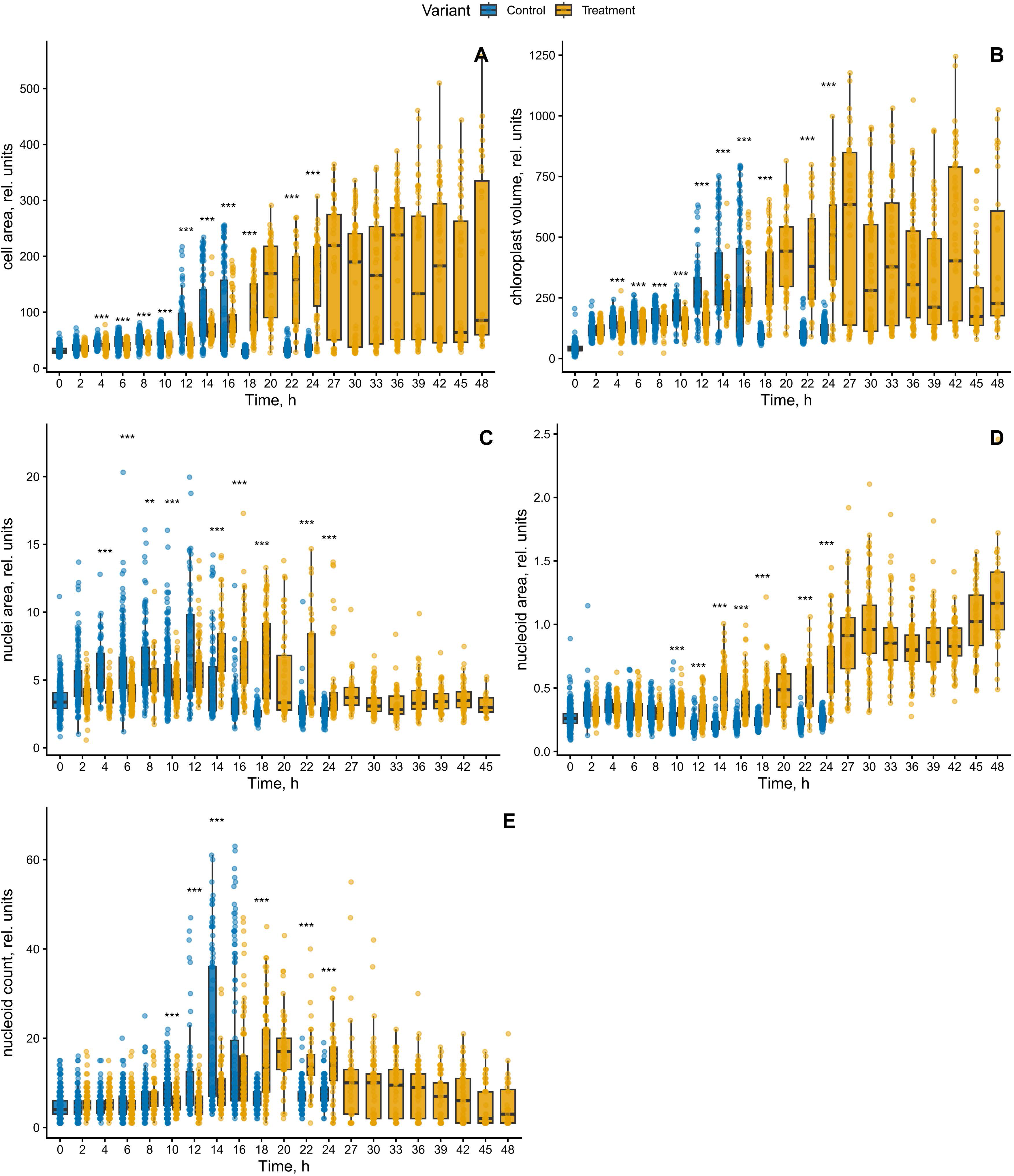
Single-cell dynamics of cellular and nucleoid parameters under control and nalidixic acid treatment in *Chlamydomonas reinhardtii*. Time-course analysis of single-cell measurements in control (blue) and nalidixic acid–treated (yellow) cultures over 0–48 h, data presented as boxplots with individual cell measurements overlaid, (A) cell area, (B) chloroplast volume, (C) nuclear area, (D) chloroplast nucleoid area, and (E) nucleoid count

**Figure 8.**
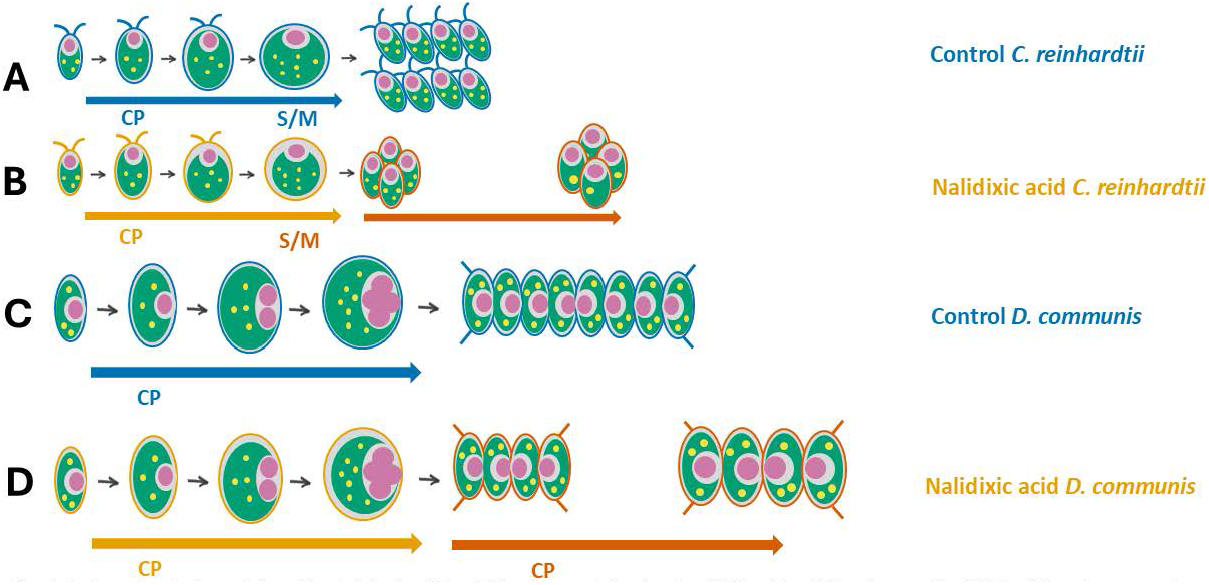
Species-specific cell-cycle organization and its perturbation by nalidixic acid. Schematic representation of growth and division patterns in *Chlamydomonas reinhardtii* (A, B) and *Desmodesmus communis* (C, D) under control (A, C) and nalidixic acid (NAL) treatment (B, D). (A) In control *C. reinhardtii*, cells grow during an extended G1 phase until they attain commitment point (CP), followed by alternating S and M phases and producing daughter cells that separate. (B) Under NAL treatment, *C. reinhardtii* exhibits delayed and less synchronized division; daughter cells fail to separate and remain in palmelloid clusters. (C) In control *D. communis*, nuclear divisions occur early and repeatedly during growth, resulting in multinucleate cells prior to division and release of daughter cells. (D) Under NAL treatment, *D. communis* maintains overall growth coordination and division structure. Arrows indicate progression through the cell cycle (CP, growth phase; S/M, DNA synthesis and mitosis).

## Discussion

Here, we aimed to clarify a possible coordination between cpDNA replication, chloroplast growth, cell growth, and the cell cycle. Our results in the control cultures confirmed the established correlation and apparent coordination among chloroplast growth, cell growth, and nuclear area. All three parameters showed tight, near-linear scaling in both model organisms (Fig. 4, 5). This aligns with established findings (Kabeya and Miyagishima, 2013; Zachleder et al., 2004; Zachleder et al., 1995) and suggests a continuous sensing-and-response system that couples biosynthetic capacity (chloroplast), cellular volume, and nuclear growth, rather than discrete, phase-restricted regulation such as checkpoints. Furthermore, the data indicated that chloroplast genome scaling is achieved primarily through modulation of nucleoid number rather than nucleoid size (Fig. 4, 5 compare C and F with G and H). This may suggest that nucleoid ploidy remains unchanged and, once replicated, the nucleoids divide. We attempted to measure the fluorescence of individual nucleoids and normalize it to fluorescent beads to directly assess individual nucleoid ploidy, but this method was not reproducible enough to allow us to draw conclusions. Nevertheless, such information is necessary to understand the dynamics of cpDNA replication and division within both control and perturbed cell cycles, and fluorescent proteins of established nucleoid components such as HBD1 (Takusagawa et al., 2021) and *hu-like protein* (Karcher et al., 2009; Kobayashi et al., 2002) would be very helpful here. At the same time, this possibly positions nucleoid number as the relevant state variable that can be sensed by the cell to coordinate downstream processes such as chloroplast growth or division readiness.

CpDNA stress induced by NAL selectively disrupted cpDNA replication and/or segregation while leaving overall cell and chloroplast growth largely unaffected during the first 24 hours of treatment (Fig. 6A, B, 7A, B). This aligns with the known action of NAL on DNA gyrase/topoisomerase activity (Robreau and Le Gal, 1974; Zachleder et al., 2004) and observed behavior in *D. communis* (Zachleder et al., 2004; Zachleder et al., 1995), suggesting that biomass accumulation and organelle expansion proceed with relatively low dependence on genome integrity, and that metabolic and structural growth programs are buffered against moderate disturbances in DNA maintenance. In contrast, the organization, copy number, and partitioning of chloroplast nucleoids were highly sensitive to replication stress (Fig. 6D, E, 7D, E). In *C. reinhardtii*, prolonged exposure to NAL reduced chloroplast growth (Fig. 7B), but growth persisted (Fig. 7A), so the proportional relationship between chloroplast and cell size progressively deteriorated under replication stress (Fig. 5B). While the nuclear area remained remarkably stable for a long time, it became effectively uncoupled from both cell and chloroplast size (Fig. 5C, D), and no nuclear division was triggered despite continued biomass accumulation. This uncoupling of growth and cell cycle progression suggests impairment of the feedback mechanism that may exist between cell and chloroplast growth (see above), leading to cell cycle arrest. Nucleoids are known to contain, in addition to DNA, various structural and regulatory proteins, components of transcription and post-transcription machinery, and other proteins (Nishimura, 2024). The nucleoid component single-stranded DNA binding protein WHIRLY has been proposed to function as a redox sensor involved in chloroplast-to-nucleus retrograde signaling (Foyer et al., 2014), making it a suitable candidate for sensing nucleoid status and possibly coordinating responses.

The most pronounced effect of NAL treatment in *C. reinhardtii* was the biphasic behavior of nucleoid parameters, characterized by changes in both nucleoid number and area. During the early stages of replication stress, cells resembled the control cells and divided their nuclei, although division was delayed (Fig. 5A-C). Soon after division, at 27 h, the nucleoid number decreased and the nucleoid area increased (Fig. 5E, D). This may represent the early phase of a phenomenon observed in cpRECA mutants and during prolonged application of NAL and novobiocin (Odahara et al., 2016). Nucleoid aggregation, which reduces their numbers and enlarges their area, thus appears to be a specific reaction to any condition affecting nucleoid DNA segregation. Interestingly, the nucleoids attempted to disperse (Kamimura et al., 2018) before aggregation, suggesting that dispersion is not dependent on completion of replication. In the absence of DNA molecule segregation, the nucleoids are possibly “pulled” back by the DNA molecule tether and aggregate. This underscores the importance of nucleoid homeostasis as an active process that requires continuous replication and segregation fidelity, as well as maintenance of nucleoid dynamics for proper function (Nishimura, 2024).

In *D. communis*, the response was much milder. Even with prolonged NAL treatment, scaling among cell, chloroplast, and nuclear compartments remained largely intact (Fig. 4A-C, 6A-C), with only modest reductions in nucleoid number and weakened correlations (Fig. 4E-H, 6D, E). However, the cells divided their nuclei into only two (Fig. 2M, N), despite significant growth and reaching the critical size required for division into four cells. This may suggest that while the *D. communis* response to NAL is more robust, the same coordination between chloroplast genome status and cell cycle entry or progression is also present. The coordination appears to depend on achieving a threshold nucleoid sufficient copy number and partitionability. When this threshold is not met, the system cannot execute the normal division program, resulting in either delayed transitions or complete cell cycle arrest. If correct, this could explain the different reactions of the two organisms to NAL. *D. communis* contains a larger pool of nucleoid DNA due to a higher number of nucleoids (compare Figs 6E and 7E) and possibly higher ploidy. This pool has a higher threshold and thus sufficient buffering capacity to respond robustly to perturbations in nucleoid replication. In contrast, the *C. reinhardtii* nucleoid DNA pool is closer to the minimum required threshold, making any response to nucleoid DNA perturbation faster and more sensitive.

Taken together, our results support a model in which chloroplast nucleoid dynamics– specifically genome copy number and its proper partitioning – act as a central integrator linking organelle genome status to cell-cycle progression. NAL-induced perturbation uncouples this integrator from downstream control, revealing that successful division requires not only sufficient biomass but also a correctly configured genetic state within the chloroplast. The differential responses of *D. communis* and *C. reinhardtii* further indicate that the robustness of this coupling may depend on the species-specific status of the nucleoid DNA. More broadly, the uncoupling of organelle expansion from nucleoid dynamics highlights nucleoid regulation as a key determinant of organelle–cell size homeostasis and suggests that plastid genome dynamics can serve as a gating mechanism for division in photosynthetic eukaryotes.

## Acknowledgements

We are obliged to the technical staff of the Laboratory of Cell Cycles of Algae for excellent technical support.

